# Virulence evolution of multiple infections with vertically and horizontally transmitted fungal species in a natural plant system

**DOI:** 10.1101/2020.09.21.307413

**Authors:** Kai Fang, Jie Zhou, Lin Chen, Yu-Xuan Li, Ai-Ling Yang, Xing-Fan Dong, Han-Bo Zhang

## Abstract

The virulence evolution of multiple infections of parasites from the same species has been modelled widely in evolution theory, and the trajectories of evolution are relevant to parasite transmission mode, as well as to parasite and host population dynamics. However, experimental studies on this topic remain scarce, particularly regarding multiple infections by different parasite species. In this study, we employed the invasive plant *Ageratina adenophora* to verify the predictions made by the model. We observed that *A. adenophora* was a highly susceptible host to phylogenetically diverse foliar pathogens with mixed vertical and horizontal transmission within leaf spots. The pathogen community structure at the leaf spot level was determined by transmission mode. Over time, the pathogen community decreased in diversity; meanwhile, the vertically transmitted pathogens exhibited decreased virulence to the host *A. adenophora*, but the horizontally transmitted pathogens exhibited increased virulence to the host. Our results demonstrate that the predictions of classical models are still valid in a complex environment. Moreover, we propose that it is very important to determine whether the primary foliar pathogen of a given plant host is relevant to seedborne fungi, as this characteristic is an important factor in understanding pathogen-host interactions.

## 1. Introduction

The traditional concept of a single pathogen infecting a single host has been widely accepted in plant pathology; however, a growing number of examples indicate that many plant diseases have been caused by multiple taxa of microbes (Lamichhane and Venturi 2015, Tollenaere et al. 2016), either in crop systems (Fitt et al. 2006) or in wild plant communities (Hersh et al. 2012, Denman et al. 2018). Therefore, it has become increasingly important to understand how multiple infections affect key components of host-pathogen interactions, including within-plant virulence and pathogen accumulation, evolutionary trajectories of pathogen populations, and disease dynamics (Tollenaere et al. 2016).

Pathogen community development of a multiple infection is often affected by priority effects, in which the order and timing of species arrival determine how species affect one another (Tollenaere et al. 2016, Bass et al. 2019). For foliar disease, fungal pathogens are believed to be commonly transmitted from airborne fungal spores (horizontal transmission) (Blakeman 2007). However, seedborne microorganisms (vertical transmission) are preferentially associated with the seedling phyllosphere (Shade et al. 2017), and is a source of inoculum for the seedling (Barret et al. 2015), and have been thought to produce strong priority effects on the microbiome assembly of seedlings, which is referred to as the primary symbiont hypothesis (PSH) (Newcombe et al. 2018). In particular, several findings have indicated that seedborne fungi are associated with pathogens (Hodgson et al. 2014). Thus, we expect that the vertical transmission of latent pathogens affects community development in multiple infections of foliar disease. Subsequently, the question becomes how virulence evolves in such a complex multiple-infection system.

Because multiple infections are common in humans and in wild systems, virulence evolution has been modelled widely in evolution theory, and evolution trajectories depend on pathogen transmission mode, as well as pathogen and host population dynamics (Alizon et al. 2013). Most models assuming that parasites are transmitted horizontally predict that the within-host competition of multiple infections favours increased virulence of parasites; as a consequence, the increased virulence benefits the transmission of parasites to other hosts, while the within-host competition excludes the weaker parasites and, in turn, causes pathogen diversity to decline (Vanbaalen and Sabelis 1995, Alizon et al. 2013). If the parasites are transmitted vertically, however, natural selection benefits parasites that are less harmful to host fitness; in the case that the parasites are transmitted both vertically and horizontally, the selection for low virulence becomes stronger when host density increases (Lipsitch et al. 1996). Lagging behind the progress of theoretical models of virulence evolution in multiple infections, however, most experimental studies have focused on infections consisting of multiple strains of the same species (Hood 2003). For plant diseases, there is still a lack of experimental evolution studies on virulence evolution in the context of multiple infections by different species with mixed transmission modes, which represents the true situation of pathogens associated with any given plant host in natural systems (Alizon et al. 2013, Tollenaere et al. 2016).

Alizon et al. (2013) proposed a three-step experimental evolution study on virulence, but the outline is simplified. Moreover, in the laboratory or greenhouse, it is difficult to perform serial passages by inoculating multiple infections of different species with distinct virulence by vertical and horizontal transmission on a host; in particular, it is not easy to compare virulence differences between initial and passaged pathogens because the evolution process commonly occurs over a relatively long time period. Thus, finding a suitable host–parasite system naturally occurring is the key step. Exotic plant introduction has been an unprecedented biogeographical experiment performed in nature, and compelling evidence indicates that evolution can occur in contemporary time spans (Callaway and Maron 2006); therefore, studying invasive systems may help to elucidate how a number of ecological and evolutionary processes operate at the same time (Lambrinos 2004). This system may be ideal for verifying the theoretical predictions regarding multiple infections and virulence evolution.

Exotic species have frequently invaded into a novel range while leaving pathogens behind; this phenomenon referred to as the enemy release hypothesis (ERH) (Keane and Crawley 2002). However, invasive plants inevitably encounter diverse local pathogens in non-native habitats (Flory and Clay 2013, Stricker et al. 2016). The mechanism by which invasive plants accumulate pathogens has also been partially explained in terms of ecology and evolution, involving spatiotemporal dynamics, host density and pathogen-host interactions (Flory and Clay 2013). For example, pathogen accumulation is positively correlated with invasive time (Mitchell et al. 2010) and the expansion of the geographical scope of hosts (Clay 1995); high host density increases the possibility of infection (Garrett and Mundt 1999). Reviews considering pathogens of invasive hosts indicate that both novel associations of invasive plants with native pathogens and cointroduction of pathogens with invasive hosts are common (Bufford et al. 2016, Dickie et al. 2017). Intuitively, virulence evolution in multiple infections with mixed transmission modes naturally occurs in this system over time but has not been fully characterized to date.

*Ageratina adenophora* (Sprengel) R. M. King and H. Robinson is a perennial herb of the Compositae family that is native to Central America but is a noxious weed in Asia, Africa, Oceania and Hawaii. This plant has invaded more than 40 countries worldwide in tropical to temperate regions, resulting in serious ecological impacts and economic losses (Poudel et al. 2019). Since the first record in China in the 1940s, the plant has been widely distributed in the provinces of Yunnan, Sichuan, Guizhou, Guangxi and Tibet, and it has continuously spread east- and northward with clear time information (Wang and Wang 2006). There is evidence that *A. adenophora* can infect fungal pathogens in the introduced ranges (Zhou et al. 2010, Poudel et al. 2019). In particular, our recent study indicates that *A. adenophora* accumulates diverse foliar pathogens from neighbours (horizontally transmitted); however, a primary pathogen (OTU515), which belongs to the family Didymellacea, does not occur on surrounding native plants (Chen et al. 2020b). Thus, we hypothesize that this pathogen may be cospread with the host by seeds (vertically transmitted) because *A. adenophora* disperses primarily by wind and water through minute asexual seeds (Wang et al. 2011). In this research, *A. adenophora* is thus an ideal model to verify the predictions made by models of multiple infections with mixed transmission modes.

## 2. Results

### 2.1 Isolation and virulence verification of leaf spot fungi

A total of 2705 fungi (451 OTUs) were isolated from 236 leaf spots, and a disease experiment screened 1149 pathogenic fungi (266 OTUs) on *A. adenophora*, phylogenetically dominated by the families Glomerellaceae, Xylariaceae and Didymellaceae (Fig. S2, Raw data). The proportion of pathogenic fungi varied across families, depending on invasive time; e.g., most strains from the family Didymellaceae were pathogens with an increased proportion over time. In contrast, most strains from the family Glomerellaceae were nonpathogens. The families Pleosporaceae and Nectriaceae were numerically small but contained high proportions of pathogens. At the OTU level, common pathogens were from the genera *Allophoma, Alternaria, Colletotrichum, Xylaria*, and *Neofusicoccum*; in particular, the primary pathogen OTU1 (*Allophoma cylindrispora*, Didymellaceae) frequently occurred in three invasive areas (Fig. 1a). In total, 212 (89.8%) leaf spots successfully harvested pathogens, with an average of 5.4 pathogens per spot being observed. Among these leaf spots, 165 (77.8%) contained > 2 pathogens of different fungal species (genotype) (Fig. 1b, Raw data); subsequently, these leaf spots were considered multiple infections and were employed to analyse the virulence evolution and community dynamics of pathogens.

**Fig. 1.**
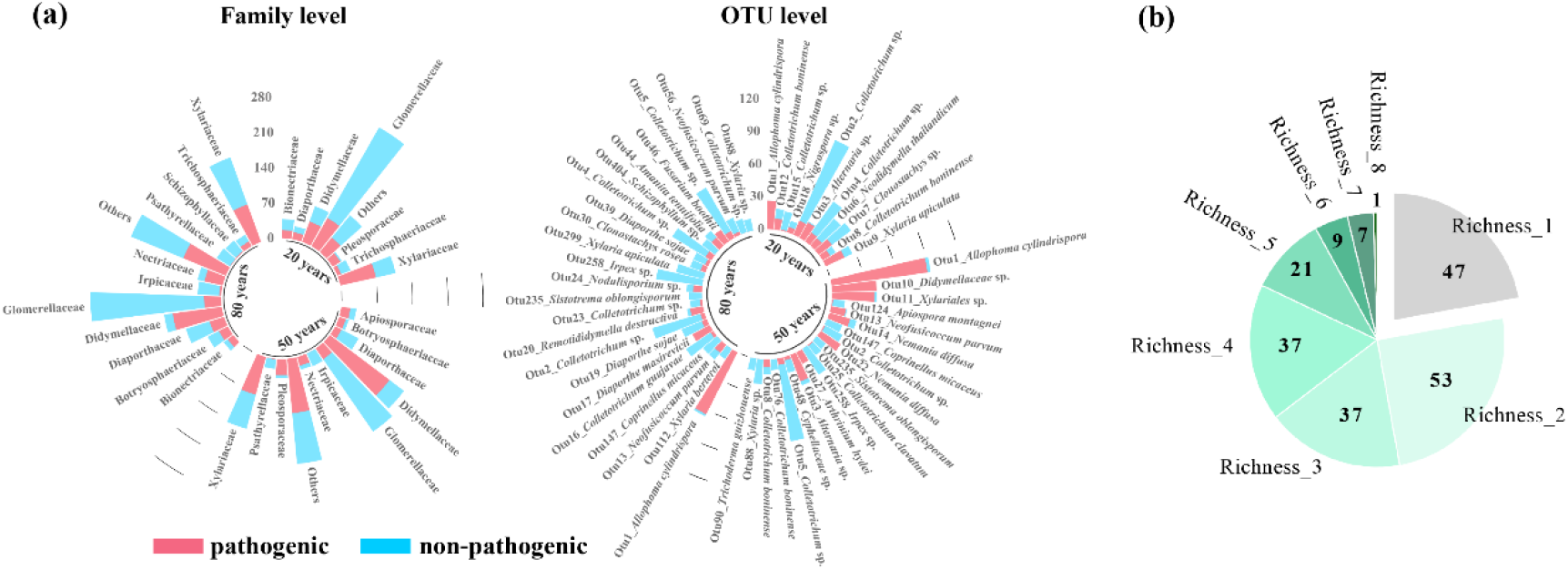
Isolation and virulence verification of leaf spot fungi. (a) The proportion of *A. adenophora* pathogens at the family (left) and OTU levels (right). Families with relative abundance > 2% are shown, and the rest were merged into “Others”; OTUs with relative abundance > 1% are shown. (b) Leaf spots with varying fungal species of pathogens. The number in each section represents the number of leaf spots with different pathogen richness. The green sections are leaf spots with multiple infections, and the grey section is leaf spot with a single infection.

### 2.2 Sharing of foliar pathogens and seedborne fungi of *A. adenophora*

The bulked seeds of nine *A. adenophora* populations were subjected to high-throughput sequencing, and 179 OTUs (591561 reads) were identified, phylogenetically belonging to 3 phyla, 27 families and 43 genera (Raw data; for rarefaction curves, see Fig. S3). By comparing the results with all isolated foliar fungi, a total of 17 overlapping OTUs were identified, accounting for 78.3% (462980 reads) and 11.5% (311 isolates) of the two fungal libraries, respectively (Fig. 2a). Among these isolates, 291 foliar isolates were pathogenic to *A. adenophora* (including 13 OTUs, accounting for 25.3% of the total fungal pathogens); the shared foliar pathogen OTU1 accounted for 21.5% of seedborne fungal reads and 15.6% of leaf spot pathogens (Fig. 2, raw data).

**Fig. 2.**
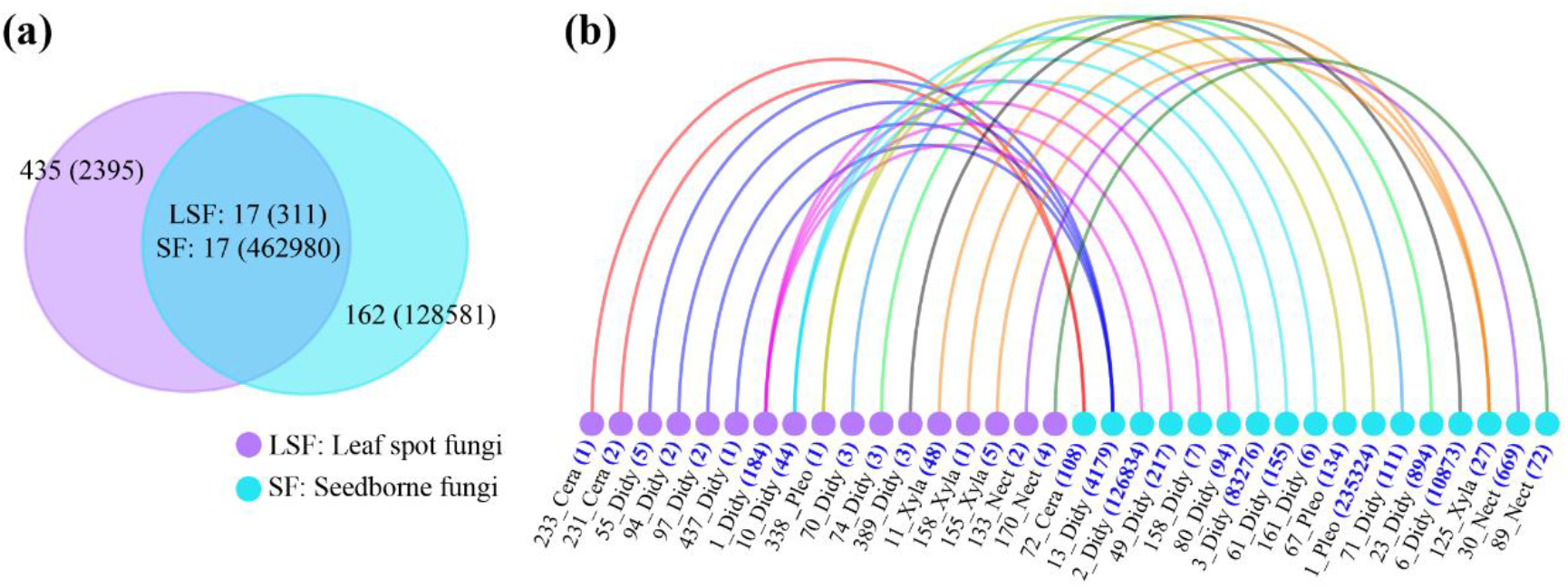
Shared OTUs between leaf spot fungi and seedborne fungi. (a) The Venn diagram shows the unique and shared OTU number (outside brackets) and abundance (reads) (inside brackets) between libraries of leaf spots and seeds. (b) The network diagram detailed the 17 shared OTUs between the leaf spots and the seeds. Didy, Didymellaceae; Xyle, Xylariales; Nect, Nectriaceae; Pleo, Pleosporaceae; Cera, Ceratobasidiaceae. The numbers in brackets represent abundance for leaf spot isolates and for seed high-throughput sequencing reads, respectively. Because the ITS sequence obtained by the high-throughput sequencing technology was short (∼250 bp), the alignment was trimmed to this range and was clustered to generate OTUs again. This trimming caused a few OTUs to merge in both libraries (see Methods).

### 2.3 Diversity and structure of fungal pathogens across invasive history at the macroscale and microscale level

At the macroscale level (geographic site), the pathogen diversity at 80 years was the highest (Fig. S4a), and pathogen communities were clustered with invasive time (F = 2.0, *p* = 0.002, 14.3% explanatory variables (EV), 7.1% adjusted explained variation (AEV) (Fig. S4b). However, at the microscale level (leaf spot) with multiple infections, pathogen diversity decreased over time (Fig. 3a). Interestingly, pathogen communities were clustered by fungal transmission mode (SP vs. non-SP) (F = 16.7, *p* = 0.002, 7.4% EV, 6.9% AEV) (Fig. 3b); 79 leaf spots contained SP pathogens, and most of them were separated from those leaf spots only containing non-SPs. In relative terms, seedborne pathogens (SPs) in leaf spots were slightly more diverse than non-seedborne pathogens (non-SPs), but both decreased with invasive time (Fig. 3c). When we pooled all 13 SP OTUs for comparison, we found that they were slightly elevated in old sites (especially at 50 years); in contrast, the non-SP OTUs were elevated in the leaf spots of newly established populations (Fig. 3d).

**Fig. 3.**
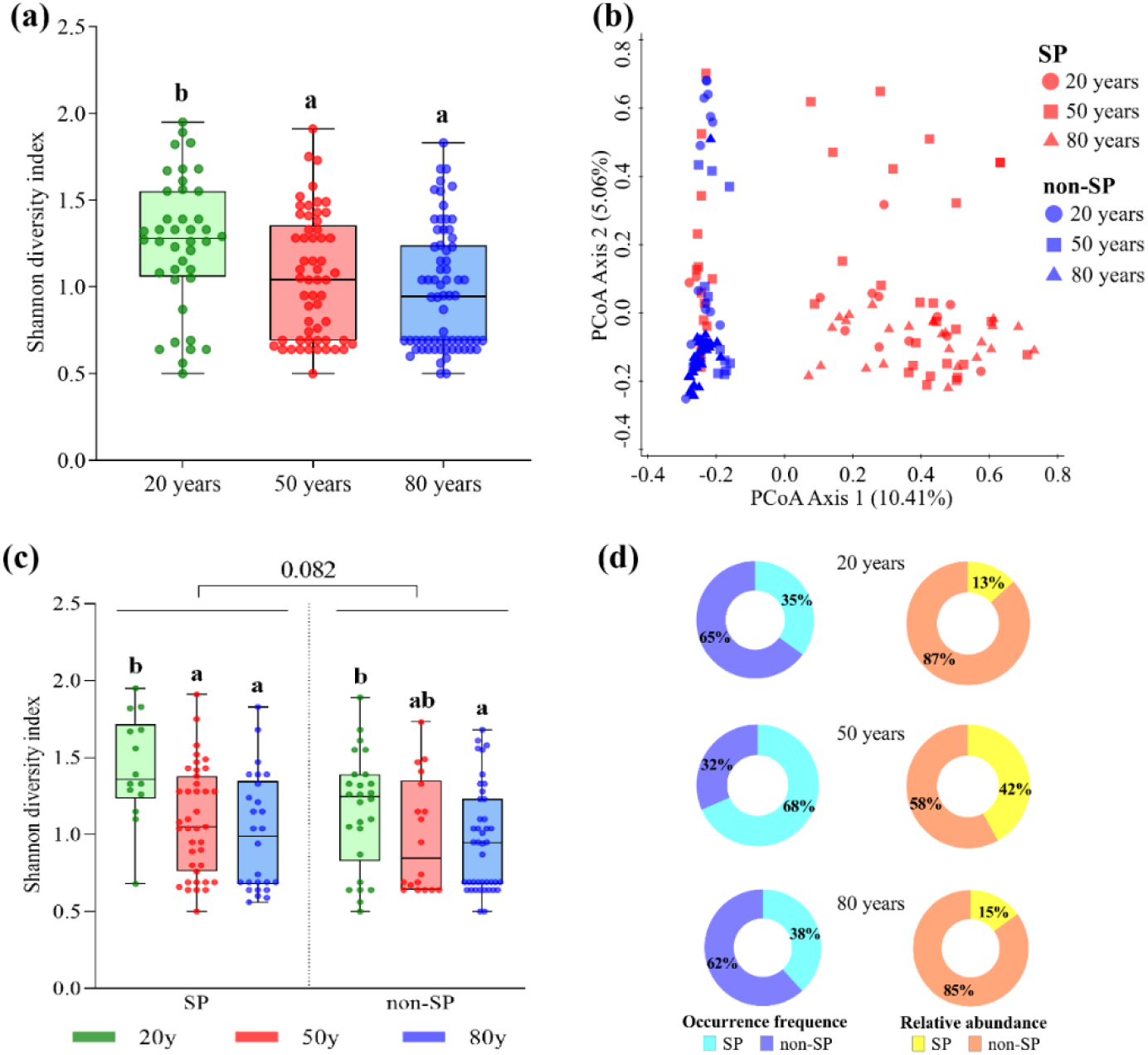
Diversity and structure of fungal pathogens across invasive history at leaf spots with multiple infections. Diversity (a) and structure (b) of fungal pathogens at the leaf spot level at different invasion times. (c) Diversity of seedborne pathogens (SPs) and non-seedborne pathogens (non-SPs) at the leaf spot level at different invasion times. Each point represents one leaf spot with multiple infections. In panels (a) and (c), nonparametric analysis with the Mann-Whitney U test was performed to show that the Shannon diversity index difference was significant among different invasion times by different lowercase letters (*p* < 0.05). In panel (b), principal coordinate analysis (PCoA) showed the similarity of pathogenic communities among leaf spots with multiple infections. SP represents leaf spot containing SP but may also cooccur with non-SPS; non-SP represents leaf spot containing only non-SPs. Percentages of total explained variation by PCoA axes in each plot are given in parentheses. (d) The occurrence frequency and relative abundance in leaf spots of SP and non-SP at different invasion times.

### 2.4 Fungal virulence evolution on host *A. adenophora*

Most of the fungal pathogens were weakly virulent to host *A. adenophora* (leaf spot area (LSA) < 50 mm^2^), accounting for 71% of the total pathogens (Fig. S5). The virulence varied within the same family, as well as within the same OTUs; however, most Glomerellaceae were weakly virulent (LSA ≤ 50 mm^2^), while Didymellaceae and Xylariaceae contained the most highly virulent strains (LSA > 200 mm^2^) (Fig. S6). We observed that the within-spot virulence variation increased linearly with pathogen diversity (Fig. 4a). On average, strains were less virulent but had higher virulence variation within spots in 20-year areas than in old areas (especially 50-year areas) (Fig. 4b). When separately analysing SPs and non-SPs, SPs showed significantly higher virulence to *A. adenophora* than non-SPs; SPs virulence decreased, but non-SPs increased, to *A. adenophora* over time (Fig. 4c).

**Fig. 4.**
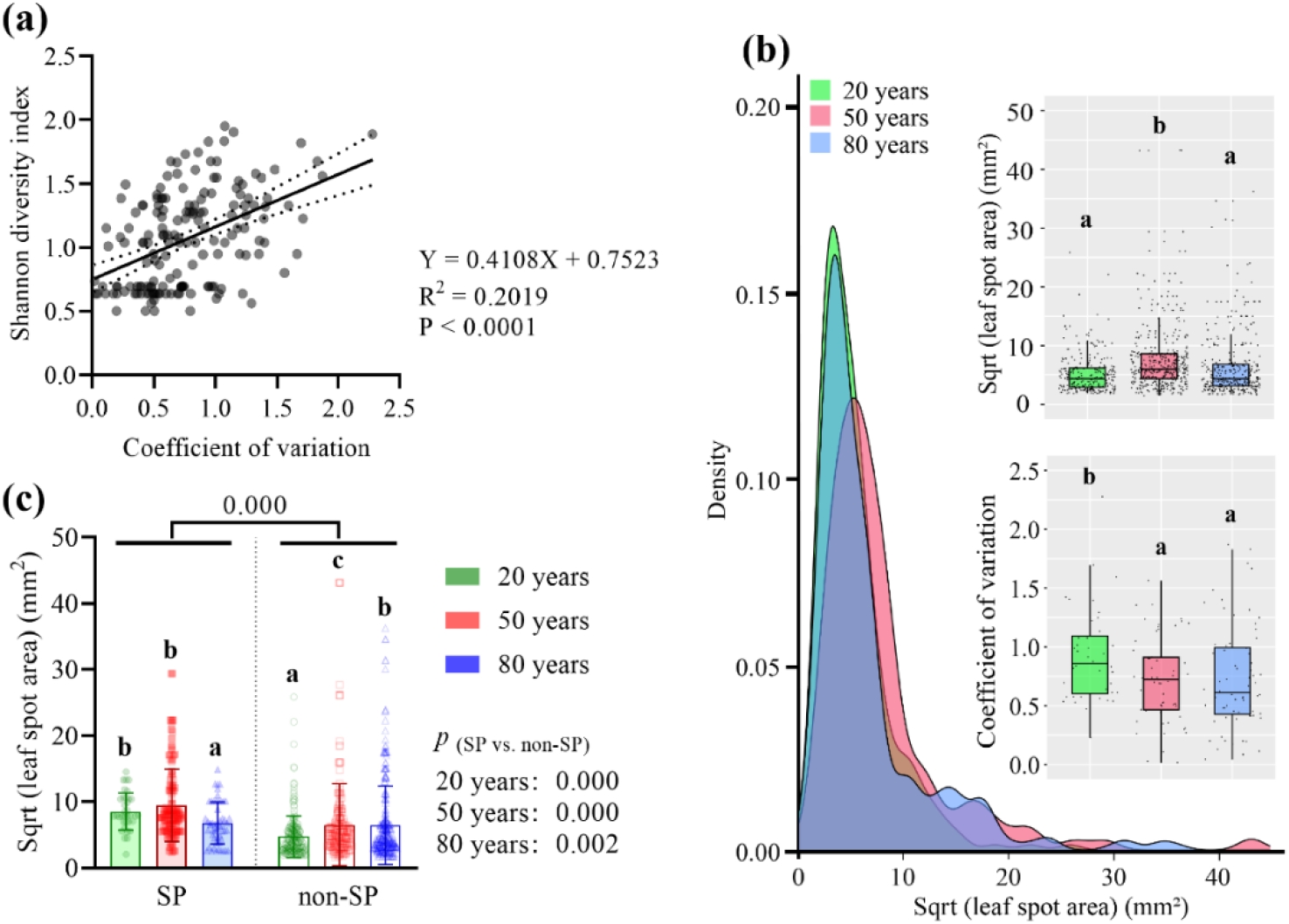
Fungal virulence evolution on host *A. adenophora*. (a) Linear analysis of within-spot species diversity and virulence variations. Each spot represents a leaf spot with multiple infections. (b) The density plots show the virulence distribution of fungal pathogens to *A. adenophora* at different invasion times. The boxplots show the differences in the average virulence (up) and the virulence variation (below) between different invasion times. (c) Virulence variation within spots of SP and non-SP at different invasion times. Nonparametric analysis with the Mann-Whitney U test was performed to test the significance of differences, and different lowercase letters indicate significant differences (*p* < 0.05). Differences between SP and non-SP of each invasion time were also given. The leaf spot area (mm^2^) is shown as square root transformed. The error bar in panel (c) represents the standard deviation.

### 2.5 Fungal virulence evolution on native plants

A total of 184 fungal pathogens (including 28 SPs and 156 non-SPs) of *A. adenophora* belonging to 88 OTUs were randomly selected to evaluate their virulence to 9 native species and to determine their host ranges. Among these pathogens, only 12 isolates were avirulent to any of the 9 native species (Fig. S7a). There were significant positive correlations among fungal pathogens’ virulence to *A. adenophora* and virulence to native species and pathogenic range (Fig. S7b). Overall, these pathogens had no change in the host range or virulence to native plants over time (Fig. S7c). In addition, both SPs and non-SPs were more virulent to *A. adenophora* than to native plants (Fig. 5a). However, SPs were less virulent to native plants and had a narrower host range than non-SPs, and this pattern also did not change over time (Fig. 5b, c).

**Fig. 5.**
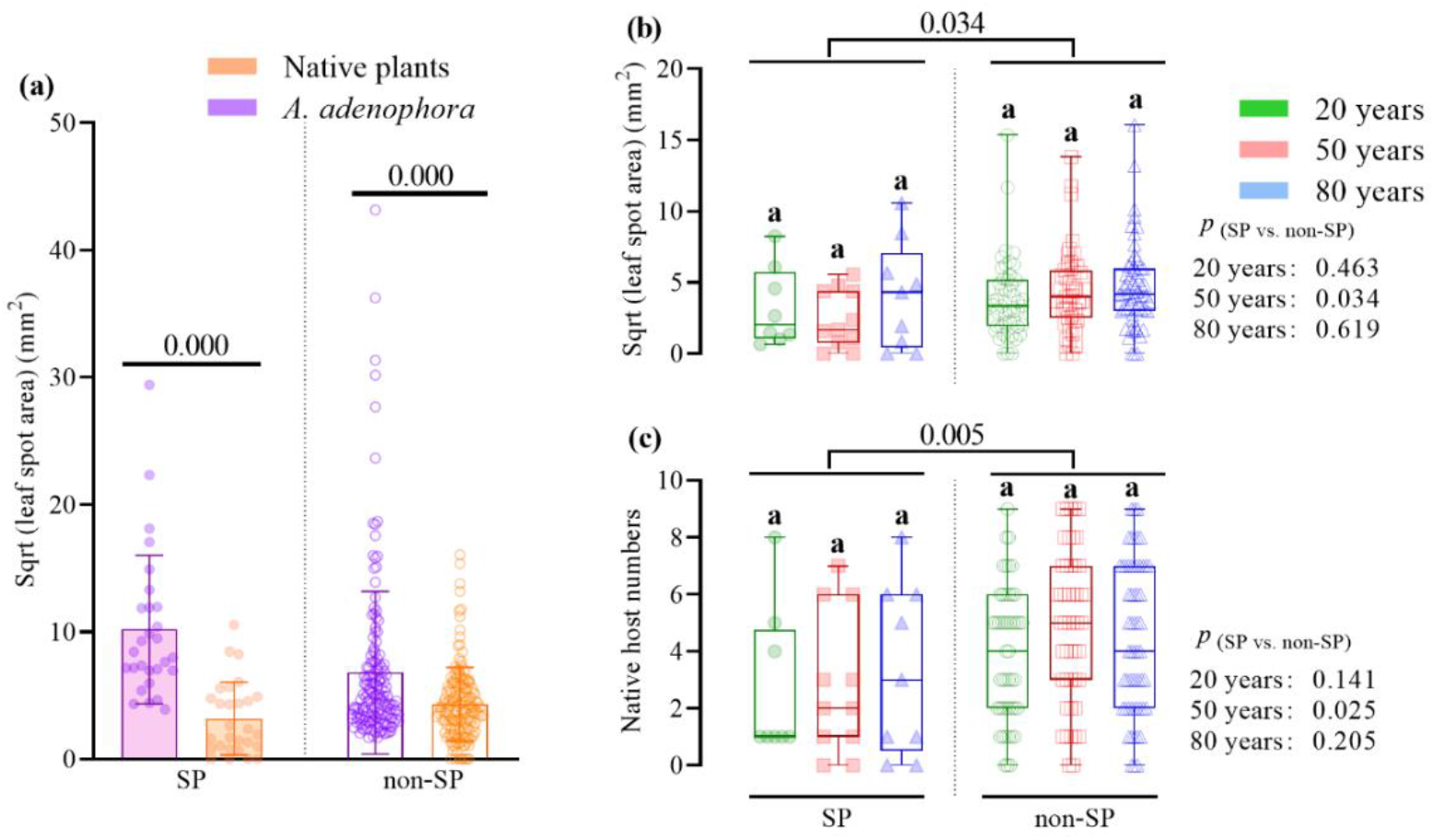
Fungal virulence to *A. adenophora* and native plants (a, b) and host range on native plants (c). Nonparametric analysis with the Mann-Whitney U test was performed to test the significance of differences, and different lowercase letters indicate significant differences (*p* < 0.05). Differences between SP and non-SP of each invasion time were also given. The error bar in panel (a) represents the standard deviation.

## 3. Discussion

Understanding how the pathogen community associated with invasive plants changes over time has been an increasingly prominent concern of invasive biologists (Mitchell et al. 2010, Bufford et al. 2016, Dickie et al. 2017). Unlike previous reports that primarily focused on the environmental, geographical and temporal patterns, as well as the ecological consequences, of pathogen emergence and accumulation (Flory and Clay 2013, Stricker et al. 2016), we employed the invasive plant system as a model to characterize the virulence evolution of multiple infections with mixed vertical and horizontal transmissions, a significant evolution theory that remains to be verified by experimental studies in plant disease (Alizon et al. 2013). Unlike many notorious invasive plants to escape enemies in the invaded range (Mitchell and Power 2003), we observed that *A. adenophora* was a highly susceptible host to phylogenetically diverse foliar pathogens in the introduced range; in particular, 77.8% of leaf spots showed multiple infections (Fig. 1b). Further experiments indicated that 25.3% of foliar pathogens were seedborne (Fig. 2a, Raw data), including one dominant pathogen, OTU1 (*Allophoma cylindrispora*, family Didymellaceae; an unidentified OTU515 was observed in our previous report (Chen et al. 2020b)), accounting for 15.6% of total foliar pathogens (Raw data).

In general, the recently introduced hosts support uniformly low pathogen richness, whereas longer-established hosts support a greater average number of pathogen species (Mitchell et al. 2010). Accordingly, *A. adenophora* leaf spot disease incidence (Fig. S1) and pathogen diversity (Fig. S4a) increased over time at the macroscale level (geographic site). However, our important finding was that at the microscale level (the leaf spot) where multiple infections are considered, this trend was reversed, with new invaders harbouring more diverse pathogens than older invaders (Fig. 3a). More interestingly, pathogen communities at the leaf spot level were clustered by pathogen transmission modes (SP vs non-SP) (Fig. 3b). The pathogen assembly of multiple infections is affected by biotic and abiotic factors (Tollenaere et al. 2016, Bass et al. 2019). Our data revealed that seedborne fungi (vertical transmission), a novel determining factor, can affect the subsequent colonization of other horizontally obtained pathogens on leaf spots. The reason for this phenomenon may be relevant to the priority effects that SPs systematically grow within hosts to produce, as the primary symbiont hypothesis (PSH) indicated (Newcombe et al. 2018). Our previous report indicated that members of the family Didymellacea (the primary SP fungi in this case, see Fig. 2) can asymptomatically grow within healthy leaves (Jiang et al. 2011) and stems (Fang et al. 2019b), and their growth is stimulated by leaf extracts (Jiang et al. 2011). That manner of growth explains why nearly 48% of leaf spots with multiple infections contain SP fungi (Fig. 3d, Raw data).

With no consideration of vertical transmission, classical theoretical models regarding multiple-strain infections from one species predict that competition exclusion within the host favours pathogenic strains with higher virulence which, in turn, causes lower pathogen diversity (May and Anderson 1983, Nowak and May 1994, Vanbaalen and Sabelis 1995). Moreover, the coinfection model predicts that increased virulence is crucial when coinfected hosts are common (Alizon et al. 2013) because increased virulence can benefit pathogen transmission between hosts to increase the exploitation rate (Hood 2003). In keeping with these predictions, we observed decreased diversity (Fig. 3c) and increased virulence (Fig. 4c) of horizontally transmitted pathogens over time. Therefore, the virulence evolution and community dynamics of those non-seedborne pathogens (non-SPs) reflect both the within-spot competition and the ecological feedback of the increased host density of *A. adenophora* over time. In the invasion front, sporadically distributed *A. adenophora* individuals encounter diverse native pathogens with low virulence due to life-history trade-offs when infecting novel hosts (Laine and Barres 2013); as more virulent strains evolve, they exclude weakly virulent strains and, in turn, cause decreased pathogen diversity over time. This process is similar to serial passage experiments (SPE), in which the initial infections with varied mutants of virulence of one species are diverse, and virulence increases as the parasite is passaged on the same host (Ebert 1998). Indeed, we observed a linear relationship between pathogen diversity and virulence variation within spots (Fig. 4a), which also implies that diverse pathogens with distinct virulence likely coinfect hosts.

In contrast to horizontally transmitted pathogens, a common notion is that vertically transmitted pathogens become avirulent because their fitness completely depends on hosts (Alizon et al. 2013). The decreased virulence of the seedborne pathogens (SPs) to the host *A. adenophora* over invasive time supported this idea (Fig. 4c). Theoretically, weakly virulent pathogens are competitively inferior in the case of mixed infections (Alizon et al. 2013). In testing this prediction, we observed that in older invasive areas, Didymellacea (primary SPs) appeared to win by increasing in abundance, as well as in occurrence frequency in leaf spots (Fig. 1a); in particular, among 47 leaf spots with a single infection, the relative abundance and occurrence frequency of SPs increased with invasive time (Fig. S8). Because most models assume that parasites are not transmitted vertically and that the evolution of increased virulence benefits their horizontal transmission to other hosts, parasites cannot increase between-host transmission rates without increasing virulence (Alizon et al. 2013). Our data indirectly suggest that vertically transmitted pathogens (SPs) can have a competitive advantage over non-SPs due to the priority effects of within-host transmission; therefore, surprisingly, the evolution of low virulence may benefit the reproduction of SPs in the mixed infections within leaf spots (Choisy and de Roode 2010).

Escaping or encountering less virulent specialists if not escaping (Reinhart et al. 2010) (Vandegrift et al. 2015) are important to plant invasions. In this case, these specialists of seedborne fungal pathogens are not easy to escape; however, their decreased virulence may not directly contribute to *A. adenophora* invasion because the foliar fungi as a whole in older sites produce stronger adverse effects on the growth of *A. adenophora* than do those in younger sites (Fang et al. 2019a). Therefore, similar the virulence evolution of non-SPs, the evolution of decreased virulence of SPs may also be in response to the increased density of *A. adenophora*, as models predict that low-virulence strains prevail, and vertical transmission becomes more prevalent, when host density increases if parasites are transmitted both vertically and horizontally on a host (Lipsitch et al. 1996). It appears that these SPs can also be horizontally transmitted, as we found a high number of Didymellacea spores in the canopy air of *A. adenophora* (Chen et al. 2020a).

A generalist strategy provides the parasite with more opportunities for transmission and persistence; therefore, in theory, decreasing specificity can be associated with increasing virulence (Barrett et al. 2009). Coinfection also offers opportunities for hybridization among different species and may lead to changes in pathogen host range (Inderbitzin et al. 2011). However, over time no evolution on native plants, including host range and virulence, occurred for both non-SPs and SPs (Fig. 5b, c), although both types of pathogen evolved in virulence to *A. adenophora* (Fig. 4c), suggesting that the virulence to the host, for fungal pathogens of leaf spots, is a more plastic trait than adaptation to novel hosts over time.

In conclusion, we employed an invasive plant as a model to answer the question of how multiple infections, particularly by different fungal species with mixed vertical and horizontal transmission mode, affect virulence evolution. Over time, the horizontally transmitted pathogens evolved increased virulence; in contrast, the vertically transmitted fungal pathogens evolved decreased virulence; meanwhile, all of these pathogens exhibited decreased species diversity (Fig. 3c). This study categorized vertically transmitted pathogens only using the sharing of seedborne fungi without verifying their within-host growth systematically one by one (see Fig. 2b). In fact, these seedborne fungi (e.g., OTU1) can also horizontally transmit (see Chen et al. (2020a)); meanwhile, some non-SPs are expected to vertically transmit if dispersal of fragments of stems and rhizomes by the flood occurs during the expansion of *A. adenophora* (Wang et al. 2011). Nonetheless, the invasive system can be treated as a virulence evolution experiment in a realistic environment; indeed, many human, animal and plant diseases exhibit mixed vertical and horizontal transmission (Lipsitch et al. 1996) and simultaneous infection by multiple species and genotypes of the same species(Lamichhane and Venturi 2015, Tollenaere et al. 2016),(Crous et al. 2009, Perez et al. 2010). Surprisingly, many predictions in classical models regarding multiple infections of strains from the same species (Alizon et al. 2013) remain valid in such a complex system, in which the primary transmission mode (vertical vs horizontal) determines the virulence evolution trajectory of pathogens in response to host density.

In addition, our finding that the assembly of foliar pathogens in leaf spots is related to seedborne fungi may have far-reaching impacts on the understanding of what factors determine the pattern of foliar pathogens in a plant community. A complex biotic (e.g., inoculum source in air and microbial interactions) and abiotic factors (e.g., microclimate) have been thought to determine the foliar pathogen community (Blakeman 2007); in particular, host plant identification strongly affects the occurrence of foliar fungal pathogens (Hönig et al. 2013, Hantsch et al. 2014). Until very recently, the primary symbiont hypothesis (PSH) proposed that a single genotype (the seedborne primary symbiont) is commonly selected due to the strong seed filter of evolution adaptation and has strong effects on the microbiome assembly of seedlings (Newcombe et al. 2018). In this study, we propose that it is very important to identify whether the primary foliar pathogen of a given plant host is relevant to seedborne fungi, which play important roles in structuring the pathogen community of multiple infections (mostly horizontally transmitted from the surroundings). In fact, the infection of a primary foliar pathogen may be common in native (Garcia-Guzman and Dirzo 2001) and invasive plants (Stricker et al. 2016), but it remains to be verified whether this pattern is relevant to seedborne fungi. On this point, we also emphasize studying pathogens at the genotype level (Callahan et al. 2016) because identifying a given genotype, rather than one species, may be crucial to elucidating the pathogen community of multiple infections.

## 4. Materials and methods

### 4.1 Study sites and sample collections

We focused our multiple infections at the leaf spot level, rather than the host individual level, because (i) a multiple infection within-leaf spot should include both direct and indirect interactions of pathogens (Tollenaere et al. 2016) and (ii) multiple fungal species or genotypes from one species have been observed to co-occur in the same spots of *Eucalyptus* (Crous et al. 2009, Perez et al. 2010); thus, multiple infections in one leaf spot should be common but remain to be characterized. Previously, we reported the foliar pathogen sharing of *A. adenophora* with local plants in Yunnan Province, which is a relatively old invasion area (Chen et al. 2020b). In this case, we expanded the sampling areas into two newly established neighbouring provinces, Guizhou and Guangxi. In a previous report, two old sites (MJ, NH) with very few fungal pathogens were excluded from analysis in this study. Moreover, because we were concerned with multiple within-leaf spot infections, pathogens without clear leaf spot sources were also excluded from the analysis. According to the geographical expansion dynamics of *A. adenophora* in Southwest China (Wang and Wang 2006), a total of 61 populations of *A. adenophora* were selected for investigation of leaf spot disease incidence, including 24, 19, and 18 populations in 80-, 50- and 20-year-old areas, respectively (Table S1). In each population, 10 individuals of *A. adenophora* were randomly selected for investigation, and the leaf spot disease incidence was calculated by dividing the number of infected leaves by the total number of leaves. We found an increase in leaf spot disease incidence with invasive time, suggesting pathogen accumulation by *A. adenophora* in the introduced range; next, at each site, leaf spots with different symptoms were collected from at least three individuals of *A. adenophora* to isolate leaf spot fungi within 12 hours after collection (Fig. S1, Raw data).

### 4.2 Isolation and identification of leaf spot fungi

In prior research, symptoms have rarely been observed on the leaves of *A. adenophora* for any of the biotrophic pathogens, such as rusts and smuts. Therefore, we focused on isolating necrotrophic pathogens. The leaves were rinsed with tap water and then surface sterilized (2% sodium hypochlorite for 30 s and 75% ethanol for 2 min and rinsed with sterile water 3 times). Healthy leaf tissues and the margins of diseased tissues of each leaf spot were cut into twelve ∼6-mm^2^ sections. For leaf spots, the size is too small (< 2 mm in diameter) to cut into 12 fragments, and 3-4 leaf spots with the same symptoms from one leaf were used to represent one leaf spot. The disinfected fragments were subsequently plated onto potato dextrose agar (PDA) and incubated at ambient temperature for 6–8 d or until mycelia were observed to grow from the leaf fragments. All fungal colonies grown from the leaf fragments were purified and used in phylogenetic analysis and disease experiments. The fungi were maintained as pure cultures at Yunnan University (Kunming, China).

Total genomic DNA was extracted from fungal mycelia using the CTAB method (Stewart and Via 1993). The ITS region was amplified with the fungal primer pair ITS4 and ITS5. Each 50-μl PCR volume included 5 μl of 10 × amplification buffer, 5 μl of dNTP mixture (2 mM), 1 μl of each primer (10 μM), 0.25 μl of Taq DNA polymerase (2 U/μl), 1 μl of template DNA, and 36.75 μl of ddH2O. The amplification was run in a Veriti 96-Well Thermal Cycler (Applied Biosystems Inc., Foster City, CA, USA) (4 min at 94 °C followed by 35 cycles of 1 min at 94 °C, 1 min at 54 °C, 1 min at 72 °C, and 10 min at 72 °C). PCR products were tested with 1% gel electrophoresis and were subsequently sent to Sangon Biotech Co., Ltd. (Shanghai, China) for ITS sequencing. Based on the GenBank and UNITE databases, sequence homology analysis, quality assessment and correction were conducted. ClustalX 2.1 was used to cut out chimeric bases to make each sequence measure approximately 550 bp in length (Thompson et al. 1997).

We grouped fungi into operational taxonomic units (OTUs) based on the internal transcribed pacer (ITS) locus because using traditional morphological species recognition and biological species recognition of fungi is operationally difficult due to the unavailability of taxonomically informative morphologies or mating systems for many of the cultured fungi (Gazis et al. 2011). A 97% cutoff for ITS sequence similarity has been demonstrated to provide a good approximation of biological species (Lindahl et al. 2013). However, determining fungal structures and functions below the species level can be very informative regarding the ecological niches, temporal dynamics, and population structure of pathogenic fungi (Callahan et al. 2016). Therefore, we analysed the fungal community using OTUs based on 100% cutoffs, which overlooks the intraspecific and intragenomic variations among ITS copies in fungi (Simon and Weiss 2008). Mothur 1.35.1 with the Navï e Bayesian classifier was used to identify the remaining sequences (database, UNITE_public_mothur_full_04.02.2020 (Abarenkov et al. 2020)) and to group the consensus sequences into operational taxonomic units (OTUs) at 100% sequence identity (Schloss et al. 2009). Because different trimming of alignment and the updated version of UNITE database were used in this analysis, which caused changes partially in the delimitation and phylogeny of OTUs that previously reported (Chen et al. 2020b), these ITS sequences were resubmitted and all nucleotide sequences reported in this study were deposited at GenBank under the accession numbers MT908377-MT908827(also see Raw data).

### 4.3 Detection of virulence and host range of leaf spot fungi

By definition, parasites cause harm to their hosts by decreasing their fitness, and this harm is usually referred to as virulence (Read 1994). We did not evaluate the virulence of pathogens on fitness by such means as modifying host fecundity, host survival or both (Alizon et al. 2013) because a large number of strains were detected. Conveniently, we used foliar damage as a virulence evaluation in this study, because on the one hand, these fungi were isolated from leaf spot, while on the other hand, foliar damage was also thought to indirectly modify fecundity and survival. We first tested the virulence of all isolated fungi on host *A. adenophora*. The inoculation experiment was performed as previously reported to test the virulence of necrotrophic pathogens in tropical forests (Gilbert and Webb 2007) with minor modifications, as detailed in a previous report (Chen et al. 2020b). Each fungal strain was inoculated into five mature and healthy leaves from a single individual of *A. adenophora* to test pathogenicity. At 1 week after inoculation, leaves were harvested and photographed, and symptomatic areas were measured. The disease experiment was performed over multiple years in the field, and environmental factors, such as moisture and temperature, were not controlled. However, all inoculation experiments were performed between June and the end of October, which is the primary growth season for plants in Kunming. Then, to further determine the virulence to native plants and host range of the verified pathogens of *A. adenophora*, a total of 184 fungal pathogens of *A. adenophora* were randomly selected, including 57, 59 and 68 strains in the vicinity of 20-, 50-, and 80-year-old invasive areas, respectively. Three native lianas (*Fallopia multiflora, Ampelopsis delavayana* and *Pueraria lobata*), three herbages (*Urena lobata, Hypoestes triflora* and *Arthraxon prionodes*) and three woody plants (*Celtis tetrandra, Cyclobalanopsis glaucoides* and *Lindera communis*) were selected for the target species to perform the pathogenicity test as above.

### 4.4 Investigation of seedborne fungi of *A. adenophora*

Because the vertical transmission of fungi in most plant seeds is imperfect and isolation frequency is commonly low (Newcombe et al. 2018), e.g., only 16 isolates are obtained from 800 seeds of *Pinus monticola* with one isolate per seed (Ganley and Newcombe 2006) (but also see Shipunov et al. (2008)), we had to perform high-throughput sequencing using bulked seed samples, rather than individual seeds. High-throughput sequencing was performed on seeds collected from 9 populations of *A. adenophora* in Yunnan Province (raw data). To identify the sharing of foliar fungi with seedborne fungi of *A. adenophora*, the representative ITS sequences from two libraries were selected to perform alignment. The ITS sequence obtained by the high-throughput sequencing technology was short (∼250 bp), and the alignment was trimmed to this range to cluster into OTUs again using MOTHUR software with a 100% similarity cutoff (Edgar 2013). This trimming caused several OTUs to merge in both libraries. Deletions/insertions were considered when comparing sequences. The OTUs that clustered from both sources were defined as shared fungi. The shared leaf spot pathogens were defined as seedborne pathogens (SPs); in contrast, the leaf spots without sharing were defined as non-seedborne pathogens (non-SPs). The next-gen data were submitted to GenBank under bioproject accession numbers PRJNA657950 (also see Raw data).

### 4.5 Data analysis

We pooled all fungi from each geographic site to treat one *A. adenophora* population to calculate the pathogen diversity at the macroscale level. At the microscale level, only the leaf spots with multiple infections were analysed, the pathogens in each leaf spot were treated as a pathogen community, and pathogen diversity was calculated; meanwhile, only the pathogens in multiple infections were used to evaluate virulence evolution and pathogen dynamics over time based on the categories of vertical and horizontal transmission mode. Here although we defined the fungal OTUs using 100% cutoffs of ITS sequence, actually we found that > 99% leaf spots with multiple infections contained >2 fungal species, with the exception that one leaf spot contained >2 genotypes from one species (see Raw data).

Nonparametric analysis with the Mann-Whitney U test was performed using SPSS v.22.0 software (SPSS Inc., Chicago, IL, USA) to compare the differences of all parameters between invasive times, including Shannon diversity, average virulence and native host range of fungal pathogens. Principal coordinate analysis (PCoA) visualized the similarity in fungal composition, and redundancy analysis (RDA) tested the correlations between fungal communities and invasion time using CANOCO v.5.0 software (Šmilauer and LepŠ 2014). All data represented fungal abundance, and distance matrices were constructed based on the Bray-Curtis dissimilarity index.

The maximum likelihood estimation was performed in 1000 replicates through rapid bootstrap analysis in MEGA 7.0 to show the phylogenetic clustering of strains (Kumar et al. 2016). The coefficient of variation (CV) was calculated by dividing the standard deviation by the mean to represent the virulence variation of fungal pathogens. Pearson’s coefficient matrix was calculated using R package “ggally” (Schloerke et al. 2014) to analyse the correlation between the virulence of fungal pathogens to *A. adenophora*, the virulence to native species and the host range on native species. Linear regression was employed to analyse the relationship between virulence variation and pathogen diversity (SPSS v.22.0).

A circular barplot was used to show the proportion of the fungal pathogen *A. adenophora* at three invasion times (R package “ggplot2” (Ginestet 2011)). The arc diagram shows the shared OTUs between leaf spots and seeds (R package “ggraph” (Pedersen 2020)). Boxplots or barplots of Shannon diversity, average virulence and virulence variation of *A. adenophora* pathogens were performed using GraphPad Prism 7 (GraphPad Software Inc., San Diego, CA, USA). A pie chart was used to show the species richness on each leaf spot, and a doughnut was used to show the occurrence frequency and relative abundance of SPs and non-SPs (GraphPad Prism 7). Density curves and bubble plots were utilized to show the virulence distribution of fungal pathogens to *A. adenophora* (R package “ggplot2” (Ginestet 2011)). A correlogram was drawn to visualize the scatter matrix and density distribution (R package “ggplot2” (Ginestet 2011)).

## Acknowledgements

The authors thank Tian Zeng, Li-Yuan Qin, Wen-Ti Zheng, and Zhi-Ping Yang at Yunnan University for help with sampling in the field and performing the disease experiment. Dr Huan-Chong Wang and Dr Tao Xu at Yunnan University assisted with the identification of plant species. This work was funded by the National Natural Science Foundation of China (grant nos. 31770585 and 31360153).

## Supplementary Information for

**Fig. S1.**
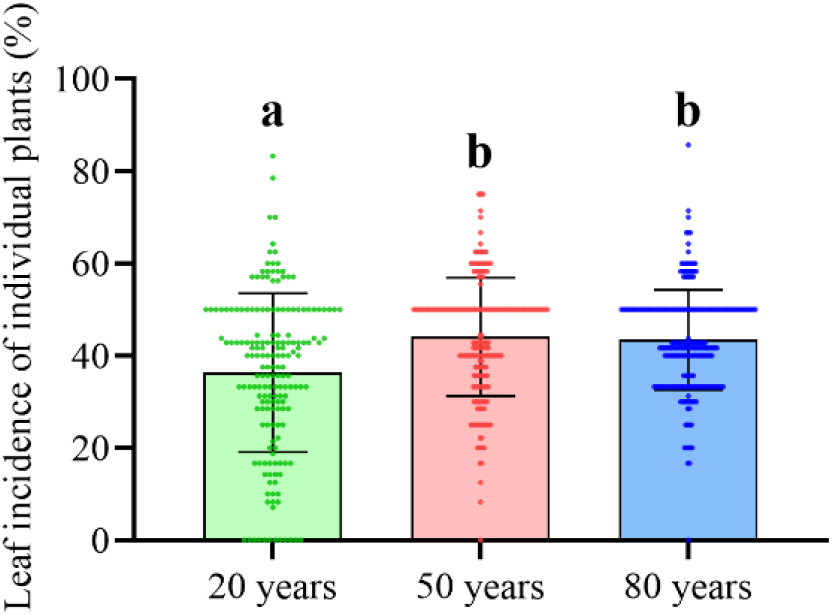
Leaf spot disease incidence of *A. Adenophora* in different invasive time. Nonparametric analysis with Mann-Whitney U test was performed to compare the differences over time, and different lowercase letters indicate significant differences (*p* < 0.05). Error bar represents standard deviation.

**Fig. S2.**
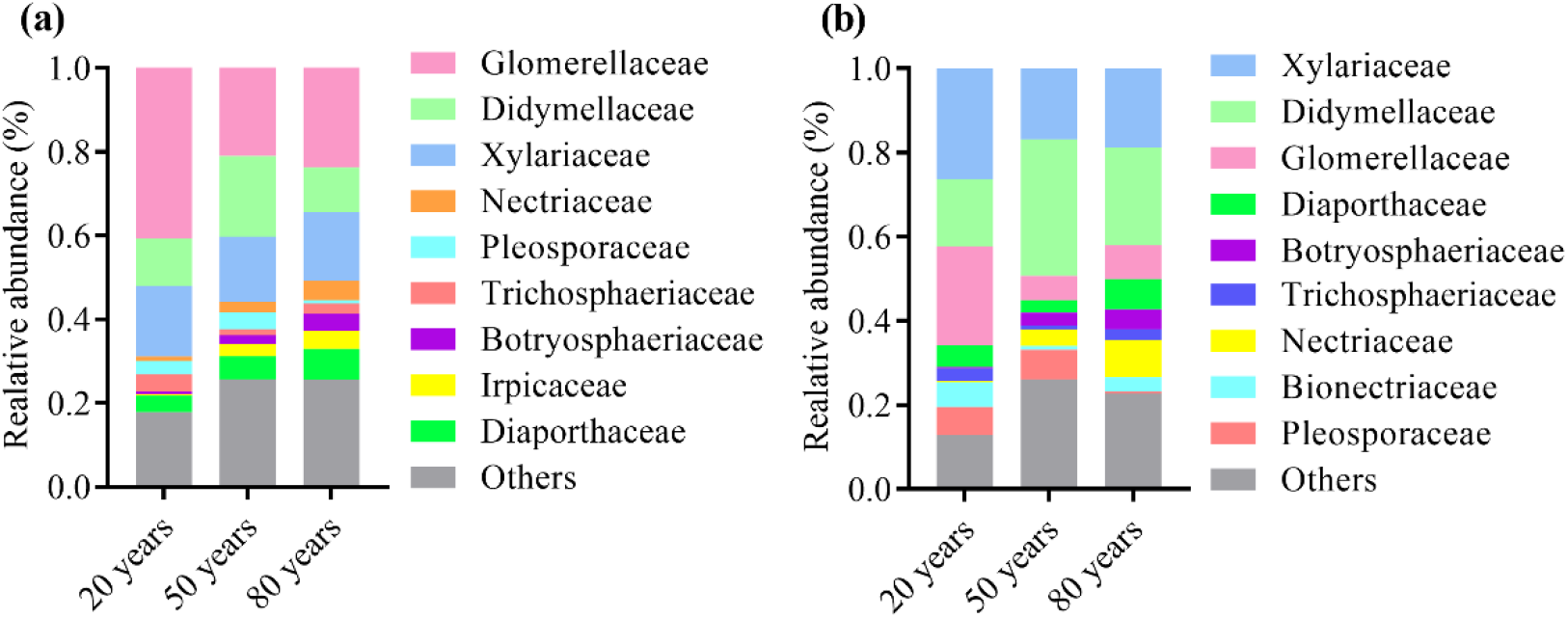
Relative abundance of numerically dominant fungi at family level. (a) Total isolated fungi. (b) The verified pathogenic fungi to *A. adenophora*.

**Fig. S3.**
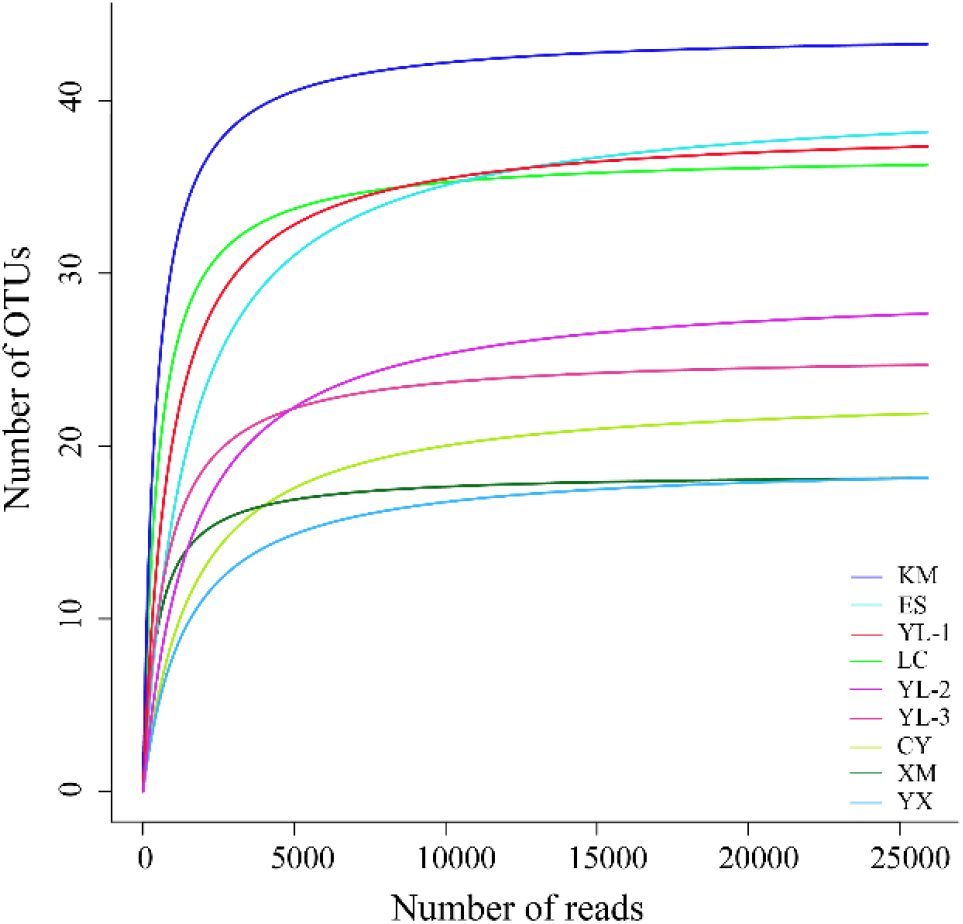
Rarefaction curves of high-through sequencing in seeds. The geographic information for *A. adenophora* population detailed in Raw data.

**Fig. S4.**
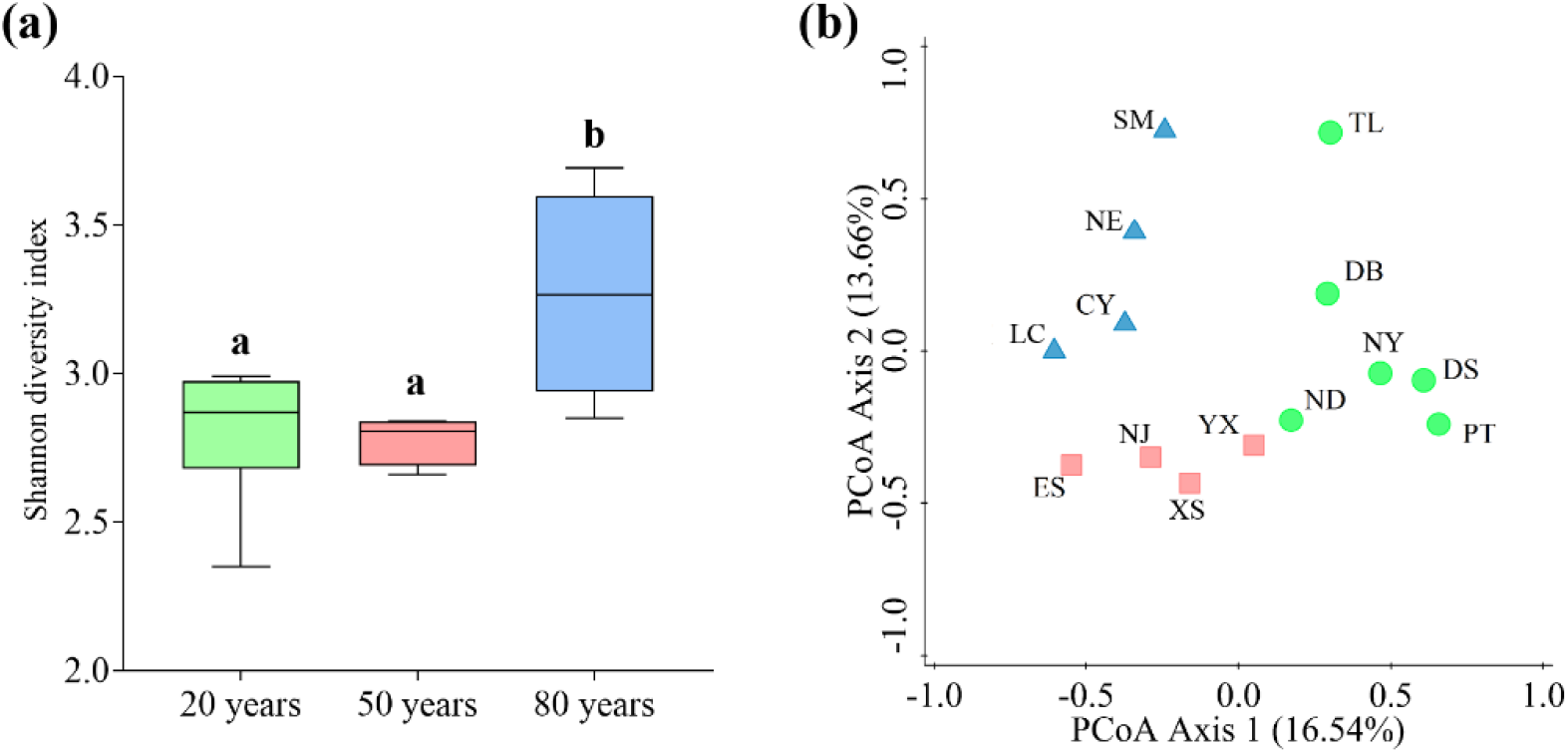
Diversity (a) and structure (b) of fungal pathogens at geographic site level. Nonparametric analysis with Mann-Whitney U test was performed to show that Shannon diversity index difference significant among invasive times by different lowercase letters (*p* < 0.05). Principal co-ordinates analysis (PCoA) showed the similarity of pathogenic fungi communities among *A. adenophora* populations. Each spot represents one geographic site (detail see Table S1). Percentages of total explained variation by PCoA axes in each plot are given in parentheses.

**Fig. S5.**
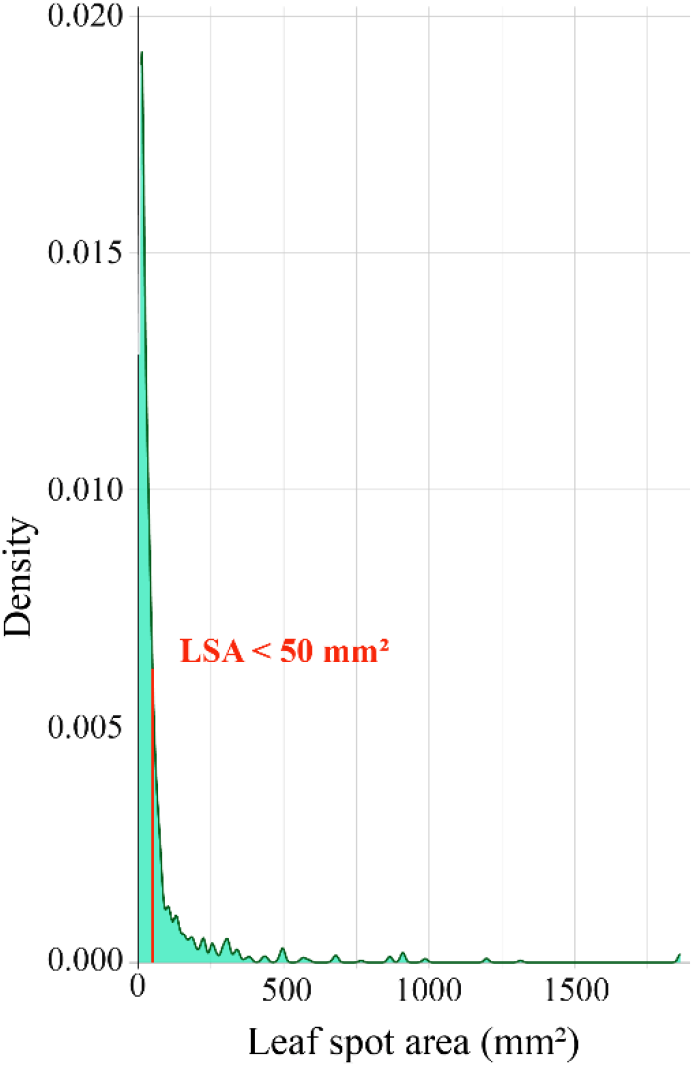
Virulence distribution of total fungal pathogens to *A. adenophora*.

**Fig. S6.**
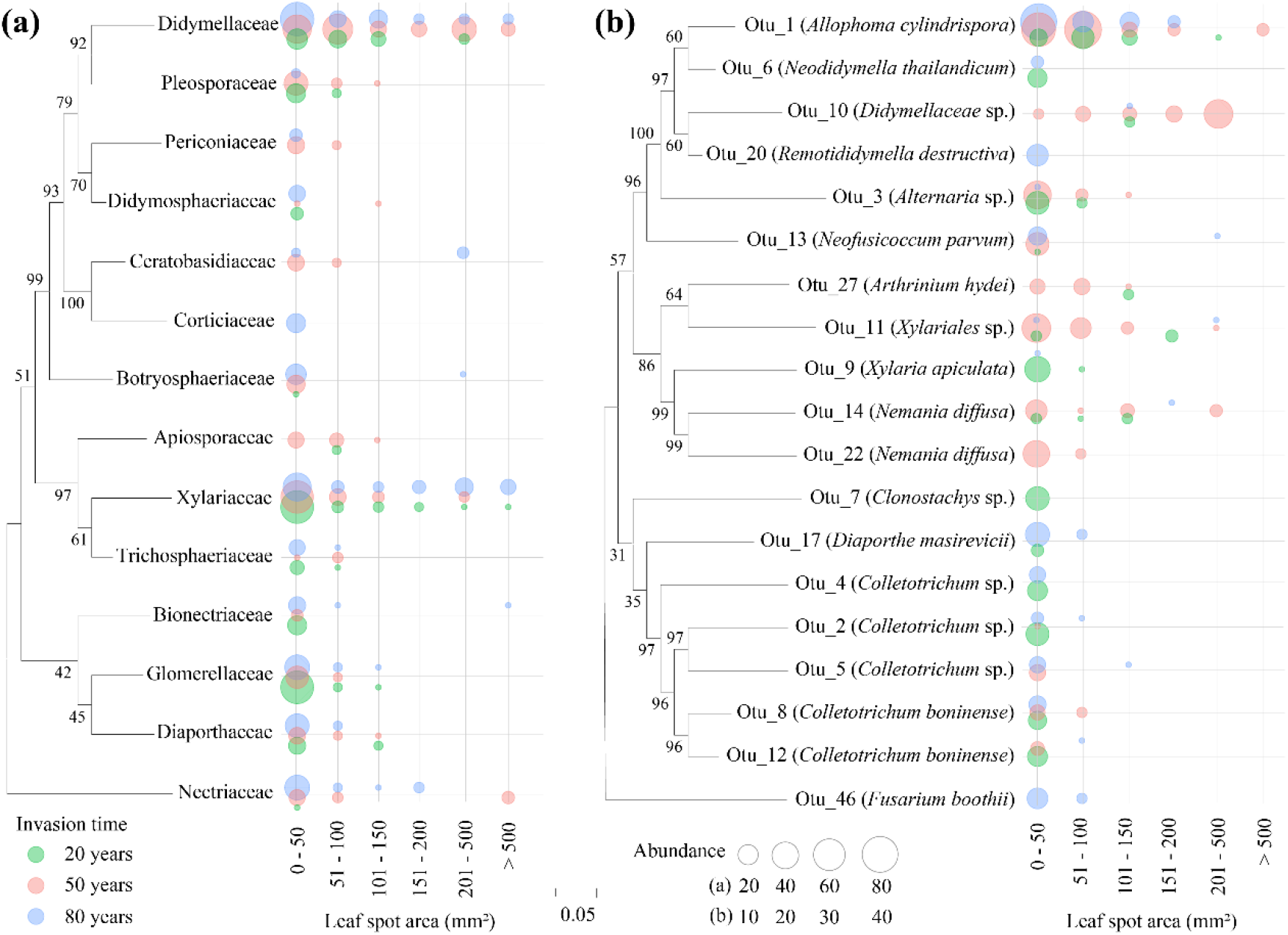
Abundance of fungal pathogens with different virulent to *A. adenophora* at family level (a) and OTU level (b). Families with relative abundance greater than 1% are shown, and the 14 relatively abundant family accounted for 86.2% of the total. OTUs with relatively abundance greater than 1% are shown, and the 19 relatively abundant OTUs accounted for 50.1% of the total. The families (or OTUs) were clustered according to phylogenetic position, and the posterior probability are indicated at the branch node. According to the density distribution of virulence of fungal pathogens to *A. Adenophora*, 5 virulence ranges (LSA ≤ 50 mm^2^, 50 < LSA ≤ 100 mm^2^, 100 < LSA ≤ 150 mm^2^, 150 < LSA ≤ 200 mm^2^, 200 < LSA ≤ 500 mm^2^ and LSA > 500 mm^2^) were selected to display the data.

**Fig. S7.**
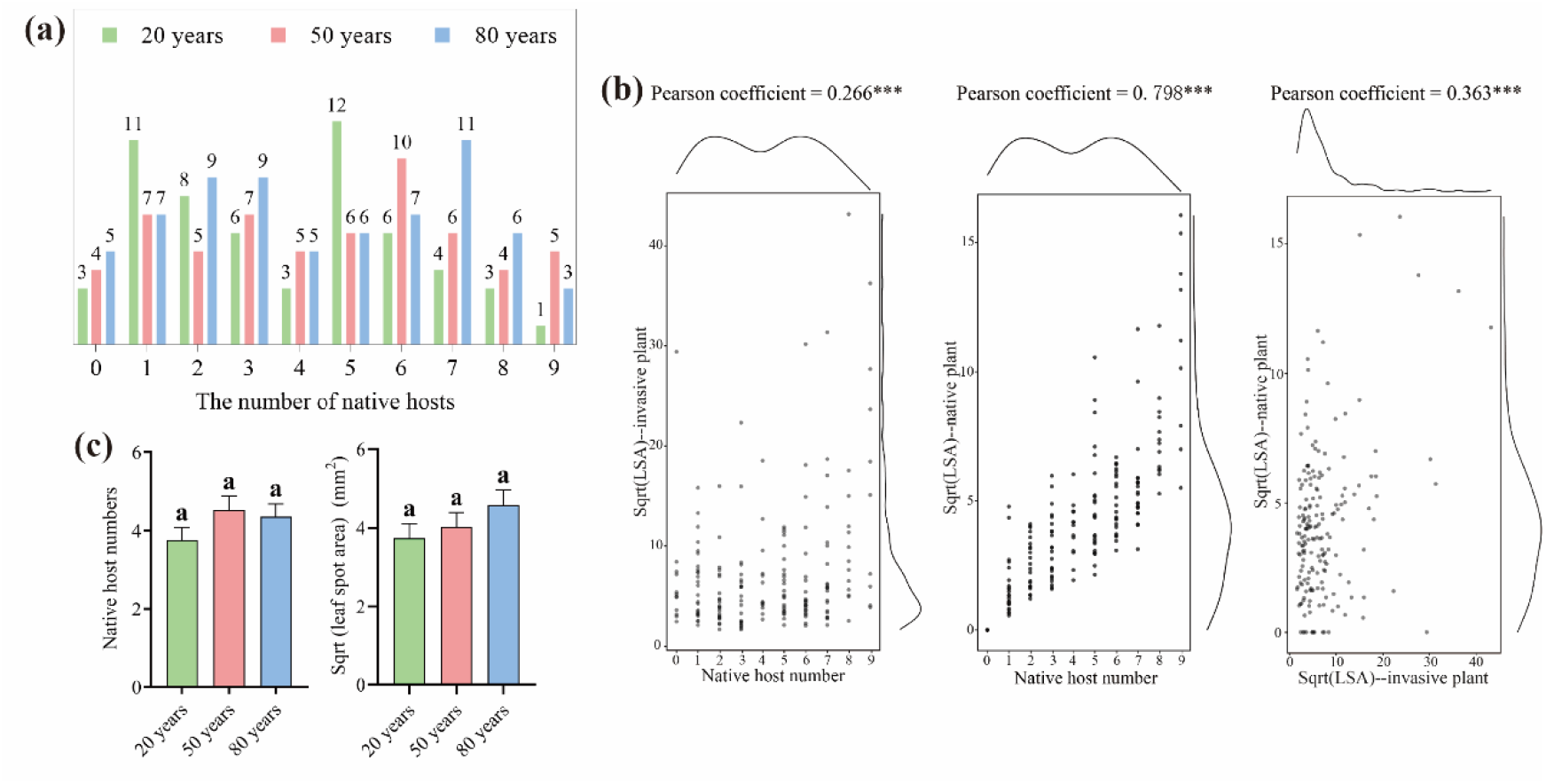
The host range and virulence of fungi to native plants. (a) Host range. The number above each bar represents the number of isolates. (b) Pearson correlations among native host number, virulence to *A. Adenophora* and average virulence to native plants of fungal pathogens. (c) Average host range and average virulence to native plants of 184 selected fungi over invasive time. Nonparametric analysis with Mann-Whitney U test was performed to test the significant of differences, and different lowercase letters indicate significant differences (*p* < 0.05). Error bar represents standard error.

**Fig. S8.**
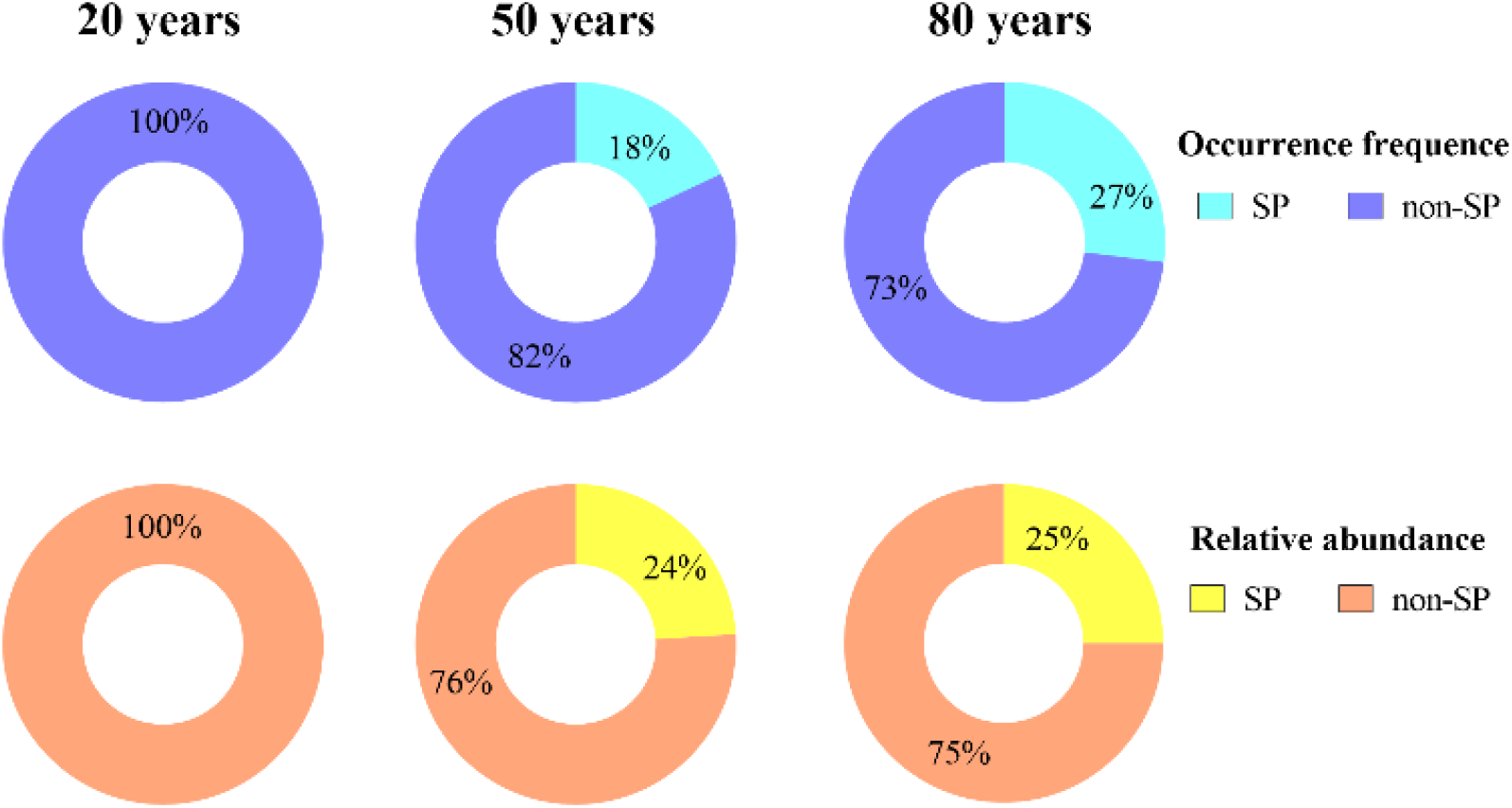
Occurrence frequency and relative abundance of SP and nonSP in leaf spots with single infection in different invasion time.

**Table S1.**
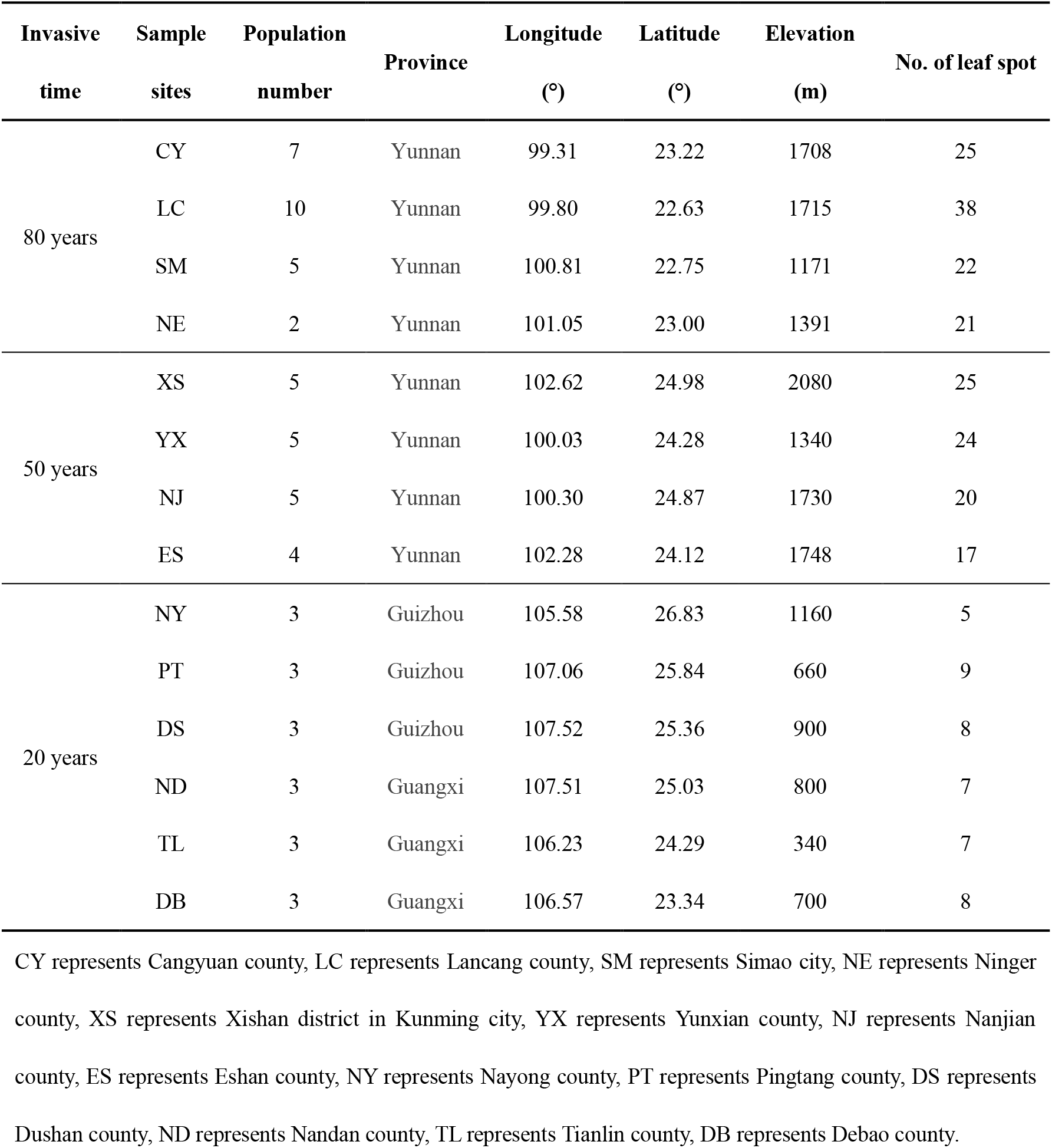
Geographical information, population distribution and number of leaf spots in three invasive times.

